# Multi-scale Modeling Toolbox for Single Neuron and Subcellular Activity under (repetitive) Transcranial Magnetic Stimulation

**DOI:** 10.1101/2020.09.23.310219

**Authors:** Sina Shirinpour, Nicholas Hananeia, James Rosado, Christos Galanis, Andreas Vlachos, Peter Jedlicka, Gillian Queisser, Alexander Opitz

## Abstract

Transcranial Magnetic Stimulation (TMS) is a non-invasive brain stimulation technique widely used in research and clinical applications. However, its mechanism of action and the neural response to TMS are still poorly understood. Multi-scale modeling can complement experimental research and provide a framework between the physical input parameters and the subcellular neural effects of TMS. At the macroscopic level, sophisticated numerical models exist to estimate the induced electric fields in whole-brain volume conductor models. However, multi-scale computational modeling approaches to predict TMS cellular and subcellular responses, crucial to understanding TMS plasticity inducing protocols, are not available so far. Here, we develop a multi-scale *Neuron Modeling for TMS* toolbox (*NeMo-TMS*) that enables researchers to easily generate accurate neuron models from morphological reconstructions, couple them to the external electric fields induced by TMS, and to simulate the cellular and subcellular responses of the neurons. Both single-pulse and rTMS protocols can be simulated and results visualized in 3D. We openly share our toolbox and provide example scripts and datasets for the user to explore. *NeMo-TMS* toolbox (https://github.com/OpitzLab/NeMo-TMS) allows researchers a previously not available level of detail and precision in realistically modeling the physical and physiological effects of TMS.

## Introduction

Transcranial Magnetic Stimulation (TMS) is a popular non-invasive brain stimulation method to safely modulate brain activity in the human brain. TMS generates a strong magnetic field by passing a transient current through a magnetic coil (Barker et al., 1985). This time-varying magnetic field crosses the skull and induces an electric field which can depolarize neurons in the underlying brain areas (Hallett, 2007). TMS is used both in research and clinical applications for neuropsychiatric and neurological disorders (Lefaucheur et al., 2014). Despite the growing use of TMS, there is still a lack of understanding of its mechanism of action.

Direct *in vivo* recordings of neural activity in rodents and non-human primates have led to key insights into TMS mechanisms (Allen et al., 2007; Li et al., 2017; Mueller et al., 2014; Romero et al., 2019). However, due to differences in brain structure and functional neuroanatomy compared to humans, great care has to be taken when translating findings across species (Alekseichuk et al., 2019). Besides *in vivo* animal studies, *in vitro* experiments in hippocampal slice cultures have been instrumental for our understanding of cellular and molecular mechanisms of TMS (Lenz et al., 2015; Tang et al., 2015; Tokay et al., 2009; Vlachos et al., 2012). *In vitro* preparations allow studying the effects of TMS on a single neuron basis in detail, however, as for animal studies, care has to be taken for translating findings to humans.

Computational modeling is a key tool to complement experimental studies to investigate TMS mechanisms. Computational models can provide a framework to understand experimental results as well as allow efficient screening of a large range of stimulation parameters. Most TMS modeling studies have focused on the spatial distribution of TMS-induced electric fields in the brain (Laakso et al., 2013; Opitz et al., 2013, 2011). These studies have been successful in predicting TMS stimulation regions and to guide TMS targeting for human experiments. However, they are limited in expanding our understanding of the TMS physiological response which depends on a variety of factors such as neuron type, electric field orientation, and ongoing activity (Di Lazzaro et al., 2018; Hannah and Rothwell, 2017). Consequently, there has been a growing interest in developing multi-scale neuron models to predict the physiological outcome of TMS.

In early modeling work, the effects of magnetic stimulation on elongated cables representing axonal tracts were studied (Basser and Roth, 1991; Nagarajan and Durand, 1996; Salvador et al., 2011). More recent work (Goodwin and Butson, 2015; Kamitani et al., 2001; Pashut et al., 2011; Seo and Jun, 2019) used sophisticated neuronal geometries. Aberra and colleagues (Aberra et al., 2020) highlighted the need to include realistic axonal reconstructions and myelination to more accurately predict neuronal responses. These studies have commonly focused on single-pulse TMS. However, for clinical applications, TMS is applied repeatedly in specific temporal patterns (repetitive TMS [rTMS]). Also, these rTMS protocols are designed to induce neural plasticity that is guided by several subcellular processes including somatic and dendritic calcium accumulation (Eilers et al., 1995; Limbäck-Stokin et al., 2004; Shoop et al., 2001). Despite the importance of rTMS-induced plasticity on intracellular calcium signaling pathways (Lenz et al., 2016, 2015; Vlachos et al., 2012), subcellular calcium-dependent processes have so far not been incorporated in computational models of TMS.

To address the limitations of available TMS models, we developed a multi-scale modeling toolbox coupling TMS electric fields with anatomically and biophysically realistic neuron models, and their intracellular calcium signaling. TMS multi-scale modeling requires the detailed knowledge of a broad range of computational tools, and so far, no easy-to-use toolboxes exist. Here, we describe a newly developed *Neuron Modeling for TMS* (*NeMo-TMS*) pipeline that allows simulating and visualizing realistic multi-scale models from neuronal reconstructions with minimal technical expertise. Our modeling toolbox allows researchers to explore TMS mechanisms computationally and embed experimental findings in a theoretical framework that can facilitate our understanding of TMS mechanisms across scales.

## Results

### Overview of Multi-scale Modeling Paradigm

We give an overview of the concept of multi-scale modeling to study the effects of TMS on neurons at the cellular and subcellular levels as shown in Figure 1. First, we use the Finite Element Method (FEM) to numerically calculate the electric fields induced in the geometry of interest (e.g. *in vitro* model or head model, Fig. 1A). However, the resulting electric fields at the macroscopic and mesoscopic scale cannot directly predict the physiological outcome. Therefore, we model the neuron membrane response to these external electric fields. To this end, we reconstruct CA1 pyramidal neurons based on microscopic images of enthorhino-hippocampal tissue cultures prepared from rodent brains (Fig. 1B). Based on the neuron morphology, we then generate a discretized numerical model of the neuron. Then, to couple the electric fields from the FEM model to the neuron model, we calculate quasipotentials (Fig. 1C) across all the neuron compartments (Wang et al., 2018). Afterward, the neuron model is numerically solved to estimate the membrane potential across the whole neuron over time (Fig. 1D). Based on the calculated voltage traces, we solve the equations governing the calcium dynamics to calculate the calcium concentrations in the neuron over time at the subcellular level (Fig. 1E).

**Figure 1.**
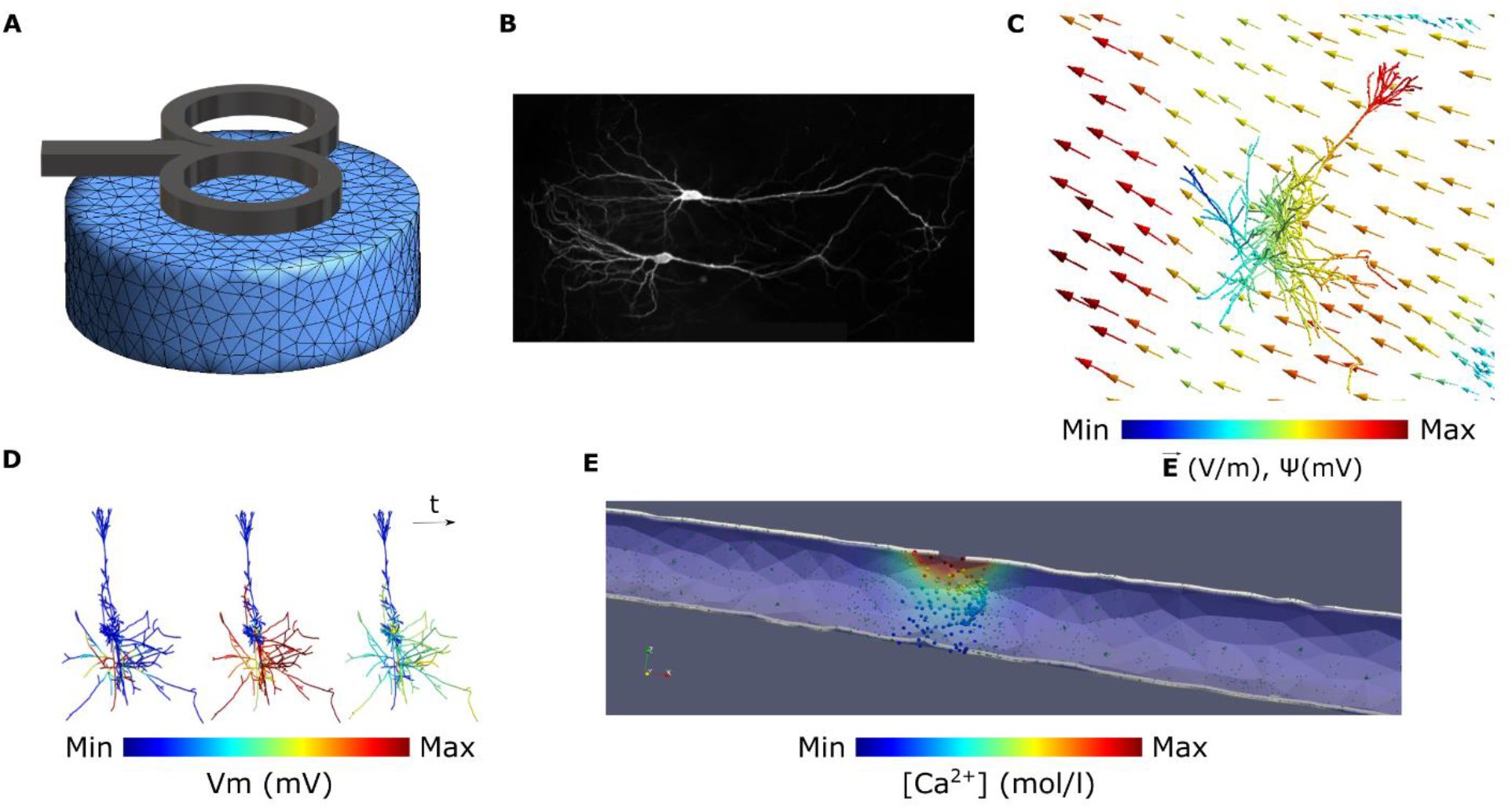
Overview of the multi-scale modeling paradigm. **(A)** Electric field calculation in the FEM model of interest. **(B)** Neuron reconstruction of CA1 pyramidal cells from microscopic images. **(C)** Coupling the electric fields 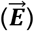 to the morphologically accurate neuron model by calculating quasipotentials (ψ). **(D)** Simulating the membrane voltage (Vm) using the quasipotentials and computing the voltage traces of the neuron compartments over time. **(E)** Simulating the release of calcium ions from the voltage-dependent calcium channels (VDCC) over time by solving the calcium diffusion equations.

### *Neuron Modeling for TMS* (*NeMo-TMS*) Toolbox

To facilitate the process of multi-scale modeling, we have developed a new toolbox (*NeMo-TMS*) and share it as an open-source resource with instructions (https://github.com/OpitzLab/NeMo-TMS) accessible to the research community. We tested the toolbox on Microsoft Window and Ubuntu. Here, we outline the toolbox functionality and the steps to perform multi-scale simulations. Furthermore, we provide examples to show how it can be used to investigate TMS-related research questions.

As shown in Figure S1, the pipeline is comprised of multiple steps that allow the user to run multi-scale models. We have shared all the necessary codes and instructions to run multi-scale models with minimal prerequisites from the user. Below we summarize typical steps in the modeling process:

1. Neuron models are generated from realistic neuron reconstructions and the biophysics of CA1 pyramidal cells are automatically added to these models.
2. Coordinates of the neuron model compartments are exported to be used in later steps.
3. The macroscopic electric fields are numerically calculated in the geometry of interest (e.g. *in vitro* model, head model). This accounts for the spatial distribution of the electric fields.
4. The electric fields computed in step 3 are coupled to the neuron model by calculating the quasipotentials at the coordinates exported in step 2.
5. Desired rTMS waveforms are generated in this step. This accounts for the temporal pattern of the electric fields.
6. The membrane voltage of the neuron is simulated based on the spatial and temporal distribution of the TMS electric fields calculated in the previous steps. Alternatively, the user can also run this step under the assumption of a spatially uniform electric field (in this case, steps 2 to 4 can be skipped).
7. The membrane voltage data are exported in formats compatible with calcium modeling.
8. The calcium concentration is simulated based on solving the calcium diffusion-reaction equations with voltage-dependent calcium channels.
9. The results from the simulations are visualized.

This toolbox is developed by utilizing multiple software packages, methods, and algorithms. Because of this and to make the toolbox accessible to a broad range of researchers with varying computational skills, we have simplified and automated the process to a great degree. For all the steps described above, the user can run the simulations using either graphical interfaces or through scripting. This feature is useful as it makes the computational workflow reproducible and gives advanced users the ability to run multiple simulations programmatically. With the *NeMo-TMS* toolbox, we provide a set of ten morphologically accurate neuron reconstructions with detailed dendritic and axonal branches to run example simulations. The morphology of these neurons is shown in Figure S2. For further technical details on the pipeline procedure, refer to the ‘methods’ section.

### Example 1: Effects of TMS on the membrane potential and calcium concentration for an *in vitro* neuron model

In this example, we run a full multi-scale simulation on an *in vitro* model and show the membrane potential and calcium activity of the neuron when a TMS pulse is delivered. As shown in Figure 2A, the *in vitro* model consists of a tissue culture placed inside a Petri dish surrounded by artificial cerebrospinal fluid (aCSF). The Petri dish is modeled as a cylinder with 30 mm in diameter and 10 mm in height. The tissue culture is 2 x 1.5 x 0.3 mm in size and is placed at the center of the Petri dish 8 mm above the bottom surface. The mesh file for this model is available for download (Alekseichuk et al., 2020). The electrical conductivity of the aCSF and the tissue culture are set to those of CSF (1.654 S/m) and grey matter (0.275 S/m) respectively (Wagner et al., 2004). A dipole-equivalent model of a Magstim 70 mm figure-8 coil (Magstim Co., UK) was placed 4 mm above the center of the Petri dish. We ran the FEM electric field simulation with a stimulator output of dI/dt = 220 A/μs. The resulting electric fields are strongest at the top center of the model since these regions are closest to the center of the TMS coil (Fig. 2B). Electric fields are aligned unidirectionally in the probe (Fig. 2C). Due to a conductivity difference between grey matter and CSF, an increase in the electric field occurs at these border walls (Opitz et al., 2015). The morphology of the reconstructed neuron is shown in Figure 2D. We placed the neuron model inside the tissue culture at the edge of the wall and oriented it in a way that the electric fields are in the direction of the neuron somatodendritic axis (Fig. 2E). Then, we coupled the electric fields to the neuron by calculating the quasipotentials across the neuron (Fig. 2F). A gradient of quasipotentials occurs in the direction of the electric field.

**Figure 2.**
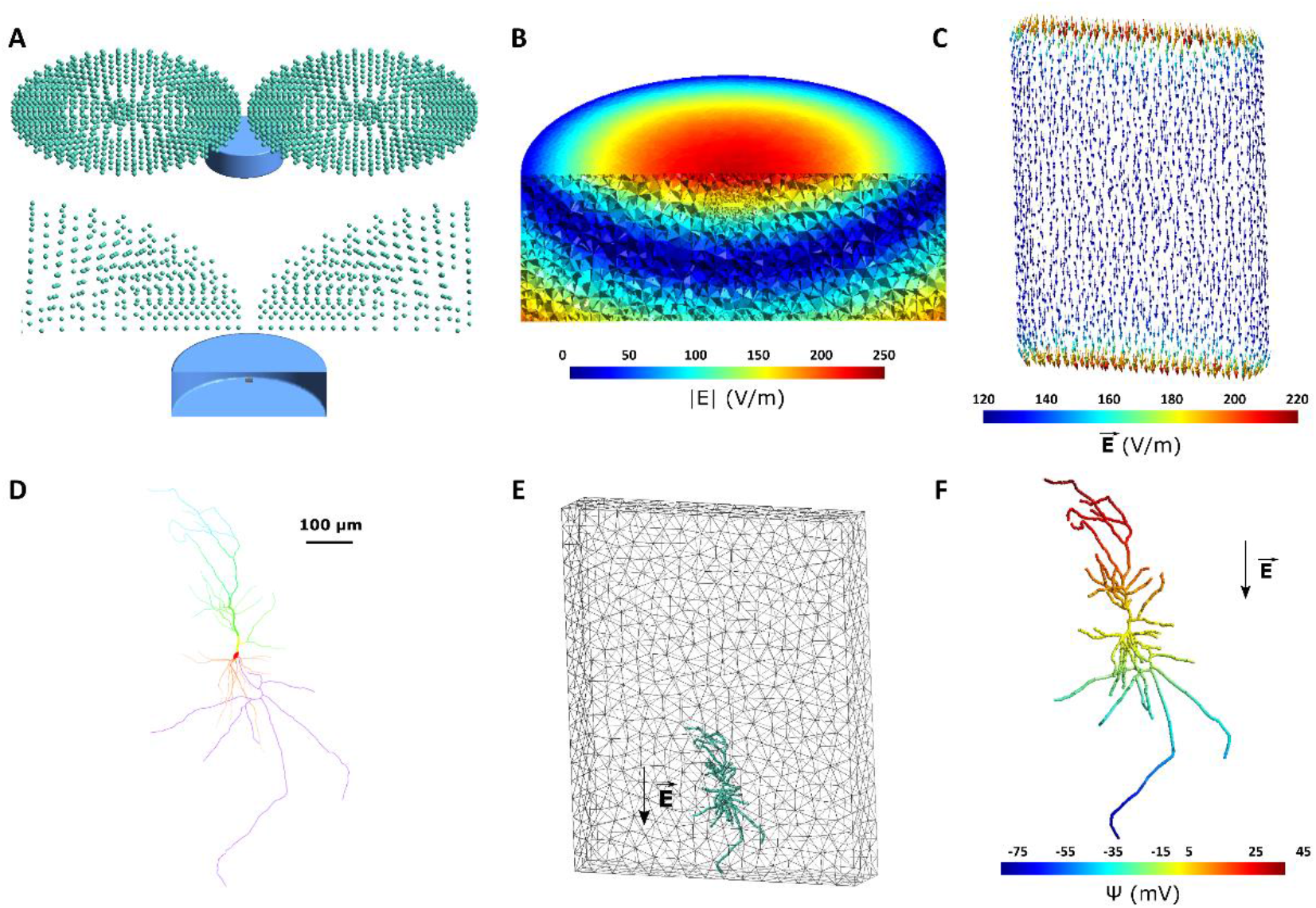
*In vitro* model of tissue culture in a Petri dish. **(A)** Geometry of the *in vitro* model. Top: TMS coil is represented through green magnetic dipoles. The Petri dish, shown in blue, is 30 mm in diameter with a height of 10 mm and is filled with aCSF. The figure-8 coil is placed 4 mm above the center of the Petri dish. Bottom: A cut-through image of the TMS coil and Petri dish is shown. The tissue culture with a size of 2 x 1.5 x 0.3 mm is placed at the center of the Petri dish 8 mm above the bottom surface. The tissue culture is modeled with grey matter conductivity. **(B)** Electric field magnitude induced in the *in vitro* model for a TMS stimulator output of dI/dt = 220 A/μs. **(C)** Electric field vector induced in the tissue culture. Electric fields are aligned unidirectionally along the handle of the figure-8 coil. Due to the conductivity mismatch between the culture and aCSF in the Petri dish, the electric field is enhanced at the borders along the electric field direction. **(D)** Reconstructed neuron morphology. Red, orange, yellow, green, blue, and purple respectively denote soma, basal dendrites, proximal apical, distal apical, apical tufts, and axon. **(E)** Neuron (green) placement inside the tissue culture (grey mesh). **(F)** The quasipotential distribution across the neuron compartments. In this model, the electric field is applied along the somatodendritic axis, thus a gradient can be seen from the apical dendrites to the axon.

Subsequently, we simulated the membrane dynamics of the neuron compartmental model using the CA1 pyramidal cell biophysics (Jarsky et al., 2005) in response to the applied electric field with the quasipotential mechanism. The resulting membrane voltage traces are then used as input to the simulation of the calcium dynamics for this neuron. While action potential initiation occurs on a millisecond timescale, calcium accumulation in the soma occurs with a delay which can be in the range of seconds in the case of rTMS. Figure 3 and the corresponding video S1 show the membrane potential of the neuron and its corresponding calcium concentrations over time during a single biphasic TMS pulse. Before the TMS pulse delivery, the neuron is at resting membrane voltage all across the cell (−70 mV). At time 0, the TMS pulse is delivered. Immediately after the TMS pulse, the axon terminal at the bottom of the cell is depolarized enough to induce an action potential. Since the axon is myelinated, the action potential quickly travels across all axonal branches and reaches the soma around 1 ms later. Afterward, the dendrites slowly depolarize as a result of ionic diffusion. Since basal dendrites are shorter, they depolarize faster than the apical dendrites. Over time (approximately 4 ms), the neuron gradually recovers back to the resting potential. Apical and tuft dendrites are the last neurites to depolarize and therefore the last ones to return to rest. The bottom panel shows the calcium densities across the neuron for the same neuron spike. Once the action potential reaches the soma at around 1 ms after the TMS pulse, with a short delay of about 0.5 ms, calcium accumulation is initiated in the soma. Then, the calcium levels start to rise slowly at the basal and apical dendrites. For these simulations, calcium exchange and release mechanisms are not considered in the axon region of the neuron; therefore, the calcium concentration remains constant in the axon of the cell. Afterward, the calcium densities in the rest of the neuron decrease and approach the resting values again (~5 ms). However, it takes longer for the calcium in the soma to fully restore to the baseline.

**Figure 3.**
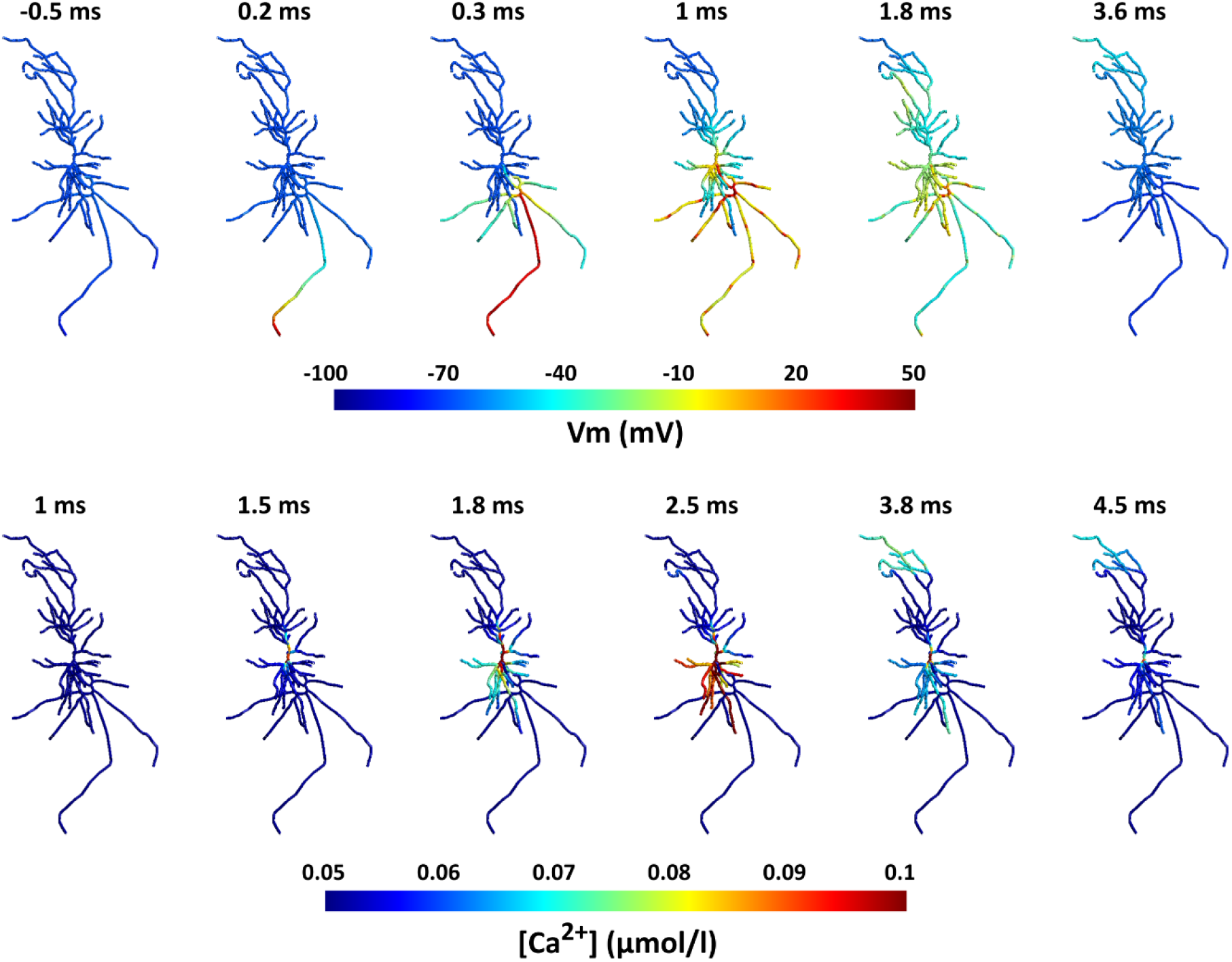
Action potential and calcium propagation over time in the neuron for the *in vitro* model. Note that time scales of membrane potentials and calcium dynamics differ between the upper and lower panel. Top: Spatial distribution of membrane potentials over time. The action potential starts at the axon terminal immediately after the TMS pulse (t = 0) and quickly propagates to the rest of the neuron. In the following ~4 ms, the neuron recovers back to its resting potential. Bottom: Distribution of the calcium concentrations displayed for the same TMS action potential. After the action potential reaches the soma ~1 ms after the TMS pulse, shortly after (~0.5 ms), the calcium concentration increases in the soma and then propagates to the dendrites. After several ms calcium levels resort to baseline. The range of the color bar for the calcium concentrations was adjusted for improved visualization and does not represent the maximum values.

### Example 2: Effect of rTMS pulse parameters on calcium dynamics

In this example, we examine the effect of rTMS pulse parameters on calcium accumulation. For this, we keep all single-pulse parameters as in example 1 and only change the rTMS protocol. We compare a 10 Hz rTMS protocol with a Theta Burst Stimulation (TBS) protocol (Huang et al., 2005). In the TBS protocol, a burst of three TMS pulses is delivered at 50 Hz repeated at 5 Hz (200 ms delay between bursts). In Figure 4, the membrane potential and the calcium concentration in the soma are shown over several TMS pulses for both protocols. After each TMS pulse, the neuron spikes, and therefore calcium accumulation in the soma follows. For the 10 Hz rTMS protocol, after each neuron spike, there is a rapid increase and then a decrease in the calcium level in the soma. However, after this initial activity, the decay rate slows dramatically. Since the calcium concentration does not completely recover to baseline before the subsequent pulse, there is a gradual increase in the overall calcium level. On the other hand, for the TBS protocol, since TMS pulses are very close together in each burst, calcium reaches higher concentrations after each burst but also decays quicker than the 10 Hz protocol. Although, because the bursts are fairly close together, the calcium level stays higher than the baseline (Fig. 4D). Overall, a buildup of calcium occurs in the soma over time in both rTMS protocols, but the temporal patterns are different.

**Figure 4.**
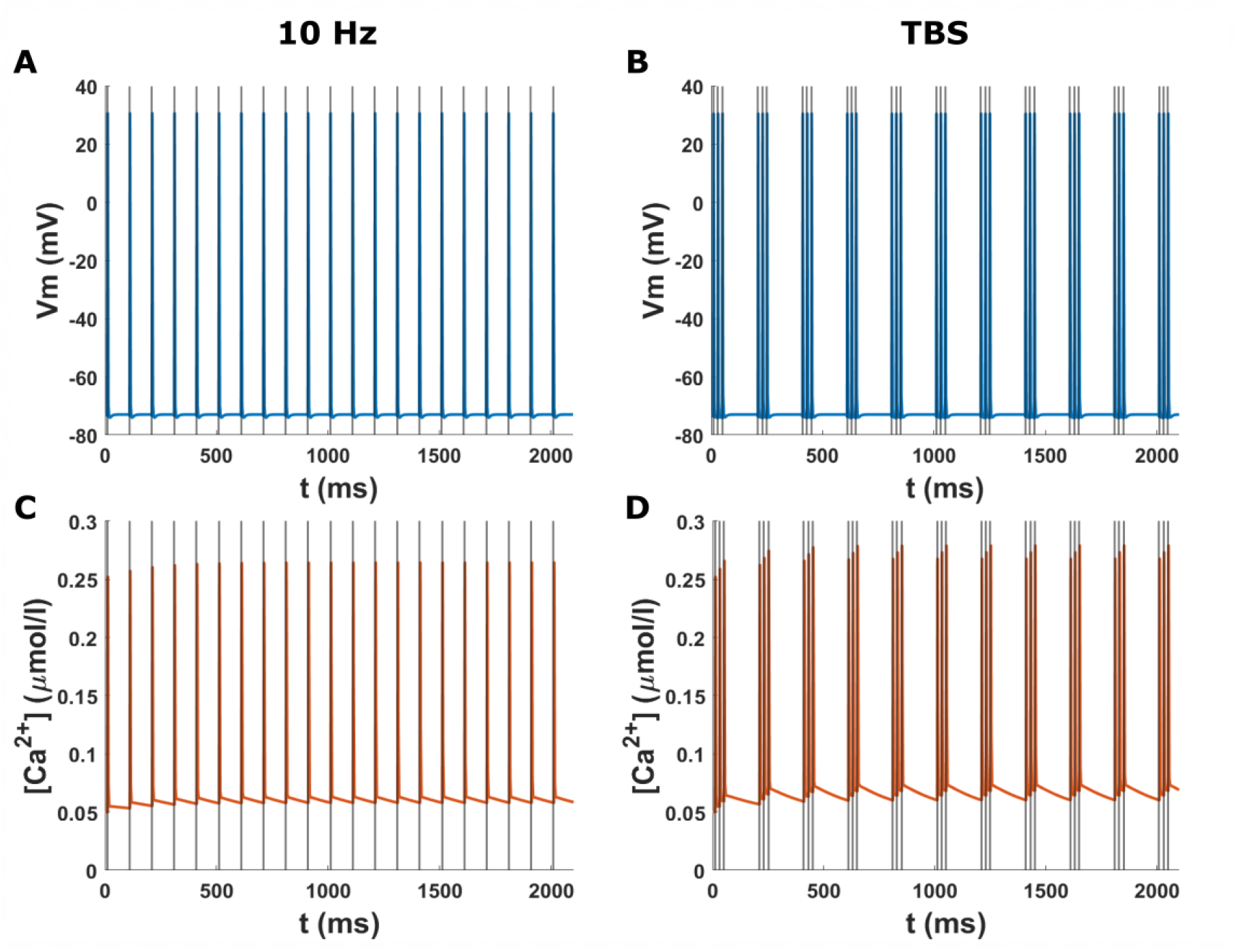
Time course of the membrane potential and calcium concentration at the soma in the *in vitro* model for two rTMS protocols. The grey lines indicate the TMS pulses. **(A)** Membrane potential at the soma for the 10 Hz biphasic rTMS protocol. The neuron spikes immediately after each TMS pulse. **(B)** Membrane potential at the soma for the TBS protocol with a biphasic TMS pulse. **(C)** Calcium concentration at the soma for the 10 Hz rTMS protocol corresponding to (A). Calcium levels rise after each spike and then slowly recover. Over time, there is a buildup of calcium. **(D)** Calcium concentration at the soma for the TBS protocol corresponding to (B). The calcium levels rise after each burst of pulses and then subside. The calcium levels stay higher than the baseline.

### Example 3: Effect of the electric field orientation on neural activation

In this example, we show how the orientation of the TMS electric field can change how it affects the neural activation site and subsequently calcium dynamics. Since the spatial distribution of the electric field plays a key role in TMS effects (Opitz et al., 2013), we compared two different electric field directions and their effects on the neuron TMS response. For this, we used one of the features of the pipeline to apply a spatially uniform electric field rather than from FEM modeling. We applied a monophasic TMS pulse in two different orientations: *i*) along the somatodendritic axis from the apical dendrite to the longest axon branch, *ii*) At 45° relative to the somatodendritic axis, along the second-longest axon branch. The neuron activation pattern is shown for these scenarios respectively in Figure 5 and video S2. In the first case, since the electric field is aligned with the long axon branch, the action potential is initiated in the terminal of the long axon branch. However, in the second scenario, the action potential is initiated in the terminal of the second-longest axon since it is more suitably aligned to the electric field. Additionally, the threshold of the electric field strength for generating the action potential differs in both cases. In the first scenario the neuron fires at an electric field strength of 170 V/m, while in the second case, a 30% higher field strength is needed for the neuron to fire. Also, there is a time shift (~0.2 ms) between the action potential initiation and propagation between these electric field orientations. This time shift causes a delay in calcium accumulation between these conditions as shown in video S3. This example shows that the electric field orientation plays a role not only in the activation thresholds but also in the neuron firing pattern, and calcium dynamics timing.

**Figure 5.**
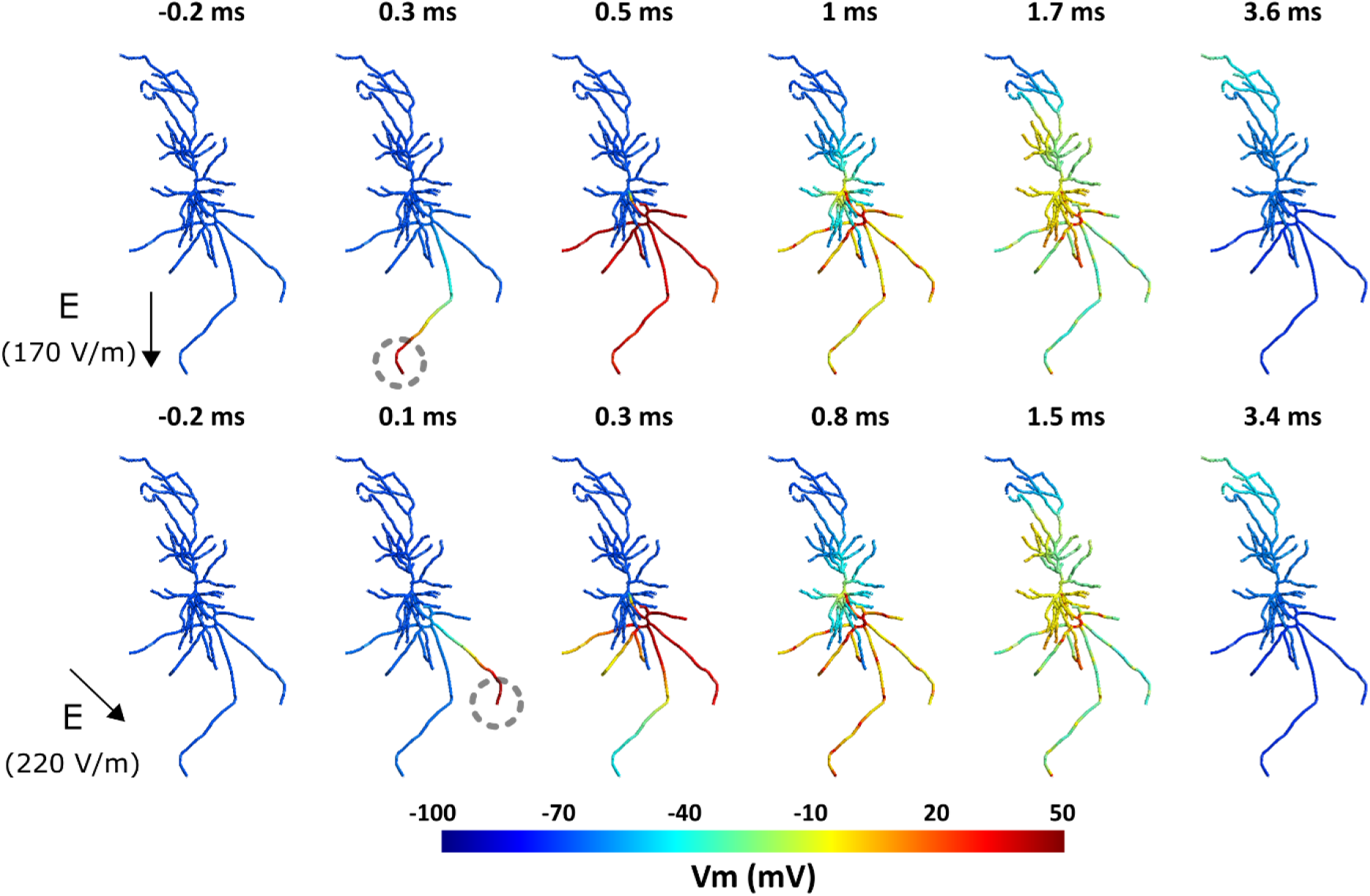
Effect of the TMS electric field orientation on membrane dynamics and spiking threshold. Top: Spiking activity in the neuron for a 170 V/m intensity uniform electric field with a monophasic TMS pulse oriented along the somatodendritic axis. The action potential initiates at the bottom-most axon terminal indicated with a grey dashed circle. Bottom: Spiking activity for a 220 V/m intensity uniform electric field with a monophasic TMS pulse oriented at 45° relative to the somatodendritic axis. The action potential starts at the axon terminal on the right.

## Discussion

We developed an open-source multi-scale modeling toolbox to enable researchers to model the effects of (r)TMS on single neurons and study their cellular and subcellular behavior. *NeMo-TMS* toolbox allows users to simulate the TMS-induced electric fields in geometries of interest (such as an *in vitro* model or a head model), to couple the TMS electric fields to morphologically accurate neuron models, and to simulate the membrane voltage and calcium concentration in the neurons. Our pipeline provides a graphical user interface, as well as an interface to run the process through scripts that will allow researchers with different computational skill sets to efficiently use our software.

To our knowledge, *NeMo-TMS* is the first modeling toolbox that enables studying single neuron behavior under TMS at macro-/mesoscopic, microscopic, and subcellular levels at the same time. Additionally, our toolbox can incorporate sophisticated neuron geometries and morphologies. Complementing modeling results with experimental studies can help to improve our understanding of the basic mechanisms of TMS.

Besides the technical implementation of the pipeline, we discuss several examples to showcase some of its capabilities. In the first example, we simulated the effect of single-pulse TMS on a morphologically reconstructed neuron embedded inside a tissue culture as an *in vitro* model. We show how the action potential is initiated at the axon terminal from which it propagates to the rest of the neuron. The voltage-dependent calcium concentrations increase after the action potential reaches the soma from which they spread into the dendrites. Both processes occur at different timescales with the calcium propagation following the action potential. In the second example, we compare the neuron response to two classical plasticity-inducing rTMS protocols: a 10 Hz rTMS protocol and a TBS protocol. We show that calcium induction varies between the protocols and that TBS results in a build-up of calcium levels. In the final example, we examine how the neuron response to TMS depends on the orientation of the electric field. For this, we applied a spatially uniform electric field at two orientations and show that the initiation site of the action potential changes as a result as well as the activation threshold. The site of the action potential initiation and the overall field intensity to initiate action potentials are in line with a recent study using morphologically accurate neuron models (Aberra et al., 2020). The differences in action potential initiation also resulted in slight delays in calcium accumulation in the soma. The exact timing between pre- and postsynaptic activity has a major impact on synaptic plasticity (Brzosko et al., 2019; Feldman, 2012; Lenz et al., 2015). It is thus conceivable that in the context of rTMS these effects may add up over the course of several hundred pulses. However, further work is required to test this prediction. Although these examples demonstrate some of the capabilities of this toolbox, its use is not limited to the examples discussed and researchers have the freedom to apply it to questions of their own interest.

While our toolbox significantly advances the field of TMS multi-scale modeling, several further developments can be envisioned. Currently, our pipeline simulates the neuron at rest without spontaneous network-driven or intrinsic activity. Additionally, neurons vary drastically in terms of their biophysics depending on their type. Here, we focused on implementing the biophysics for CA1 pyramidal neurons. Currently, the calcium simulations do not take into account internal calcium stores from the endoplasmic reticulum (ER) and only simulate the calcium release from voltage-dependent calcium channels (VDCCs), sodium-calcium exchangers (NCX), and plasma membrane Ca^2+^ ATPase (PMCA). Future versions of our pipeline can incorporate intrinsic synaptic activity, provide biophysics for more diverse neurons such as cortical neurons, and allow users to define their own biophysics. Further developments can also be implemented to incorporate modeling of the calcium in the ER. Additionally, our toolbox can be expanded to include other non-invasive or invasive brain stimulation techniques such as transcranial Alternating Current Stimulation (tACS), transcranial Direct Current Stimulation (tDCS), or Deep Brain Stimulation (DBS) in the future. Another promising avenue for future developments is modeling the effects of brain stimulation on a network of neurons. One way to achieve this is by combining *NeMo-TMS* with other neuron network modeling frameworks such as the human neocortical neurosolver (Neymotin et al., 2020).

In conclusion, *NeMo-TMS* is a unique tool that provides an easy-to-use platform for multi-scale TMS modeling and enables researchers to incorporate sophisticated modeling approaches into their research.

## Materials and Methods

### Neuron Reconstructions

#### Ethics Statement

Animals were maintained in a 12 h light/dark cycle with food and water available ad libitum. Every effort was made to minimize distress and pain in animals. All experimental procedures were performed according to German animal welfare legislation and approved by the local animal welfare officer of Freiburg University.

#### Preparation of Organotypic Tissue Cultures

Enthorhino-hippocampal tissue cultures were prepared at postnatal days 4-5 from Wistar rats of either sex as described previously (Lenz et al., 2016).

#### Neuronal Filling and Imaging

Single CA1 pyramidal neurons were identified under a microscope (LN Scope; Luigs and Neumann) equipped with a 40X objective (NA 0.8; Olympus) and a Dodt-Gradient-Contrast system. The bath solution contained 126 mM NaCl, 2.5 mM KCl, 26 mM NaHCO3, 1.25 mM NaH2PO4, 2 mM CaCl2, 2 mM MgCl2 and 10 mM glucose and was saturated with 95 % O2 / 5 % CO2. The cells were patched using 3-6 MΩ patch pipettes pulled from borosilicate glass and were filled with an intracellular solution containing 126 mM K-gluconate, 4 mM KCl, 4 mM ATP-Mg, 0.3 mM GTP-Na2, 10 mM PO-Creatine, 10 mM HEPES, and 0.1% Biocytin (pH = 7.25 with KOH, 290 mOsm with sucrose). The cells were held at −60 mV and the whole-cell configuration was maintained for at least 10 min to ensure complete filling of the cells, even at the distal dendrites. Patch pipettes were retracted carefully to allow for the cell membrane to close again and the tissue cultures were fixed in a solution of 4 % PFA (w/v) and 4 % (w/v) sucrose in 0.01 M PBS for 1 h. The cultures then were incubated for 1 h with 10 % (v/v) NGS and 0.5 % (v/v) Triton X-100 in 0.01 M PBS and subsequently for 4 h with Alexa-488 conjugated Streptavidin (1:1000; in 0.01 M PBS with 10 % NGS and 0.1 % Triton X-100) and DAPI was used to visualize cytoarchitecture (1:5000; in 0.01 M PBS for 15 min). Tissue cultures were washed with 0.01 M PBS and mounted onto glass slides for visualization with an anti-fading mounting medium. Confocal images were acquired using a Nikon Eclipse C1si laser scanning microscope with a 40x (NA 1.30; Nikon) objective. Images were acquired as multiple Z-stacks with a step size of 0.5 μm (voxel size x and y = 0.3784 μm) in a tile-scan configuration and stitched together using the FIJI software (Schindelin et al., 2012).

#### Neuronal Reconstructions

CA1 pyramidal cells were reconstructed using Neurolucida 360 (ver. 2019.1.3; MBF Bioscience). Confocal images were imported in the Neurolucida 360 mainframe as an image stack. Somata were reconstructed using manual contour tracing, with the contour tracing set to ‘Cell Body’. Dendrites were subsequently reconstructed in the Neurolucida 3D environment under the ‘User-guided’ tracing option using the ‘Directional Kernels’ method. These reconstructed cells have both detailed axonal and dendritic branching. The raw reconstructed morphological data was then imported into the TREES toolbox for additional processing. (Cuntz et al., 2010) To correct for diameter overestimation due to fluorescence halo, a quadratic diameter taper algorithm (Cuntz et al., 2007) was applied across the dendritic arbor, with separate consideration for the basal dendrites, apical tuft, apical oblique projections, and primary apical dendrite. Parameters for the diameter tapering algorithm were adapted from (Lenz et al., 2015), who estimated them based on data from (Golding et al., 2005). Internodal segments of the axon were assigned a fixed diameter of 1μm for and 0.8μm for nodes of Ranvier. As abrupt changes in the direction of a neurite cause anomalous local electric fields, a smoothing algorithm was also applied to the neurites. Using ProMesh4 (Goethe-Universität, Germany), we applied a Laplacian smoothing to all neurites (alpha = 0.25, 20 iterations) as well as manually removing any remaining anomalous sharp direction changes. These ten neuron reconstructions are shared with the toolbox as samples.

### Neuron Model Generation

We integrated a series of software tools into an automated pipeline for generating NEURON compartmental models (Hines and Carnevale, 1997) for modeling the effect of TMS on single brain cells. This pipeline is capable of generating models from commonly used file formats, i.e., SWC and Neurolucida ASCII files. Note that it is up to the user to ensure the input morphologies are correct, high-quality and without artifacts, otherwise the model generation may fail in the process or the simulation results would not be reliable. We tested the pipeline on the ten reconstructions of rat CA1 pyramidal cells provided here, as well as other morphology files.

Since the axonal reconstructions do not include myelination, this pipeline allows the user to myelinate the axon automatically, or to leave the neuron unmyelinated. For this, we implemented a modified variant of the myelination algorithm used in (Aberra et al., 2018). Nodes of Ranvier were placed at all bifurcation points in the axon arbor, as well as regularly at 100μm intervals. All internodal segments except terminal segments shorter than 20μm were myelinated. As most publicly available reconstructions of CA1 pyramidal neurons do not have an axon, the pipeline also features a provision for potential automatic addition of a straight artificial axon; in this case, the axon is a straight line emanating from the basal region of the soma with the first 10um a hillock segment, the next 15μm the axon initial segment, followed by six 100μm long myelinated internodal segments with regularly spaced 1μm long nodes of Ranvier.

#### Biophysics

The NEURON compartmental models were generated using the T2N extension of the TREES Toolbox (Beining et al., 2017), which translates the TREES Toolbox morphological data into NEURON’s HOC format and endows the model with biophysics in the MATLAB environment (Mathworks, Inc., Natick, MA, USA). Our models implement a generalized version of the Jarsky model of the CA1 pyramidal cell (Jarsky et al., 2005). This includes the passive properties: C_m_ = 0.75 μF/cm^2^, R_a_ = 200 Ω-cm, Rm = 40000 Ω/cm^2^. Additionally, axon myelinated segments had a significantly reduced C_m_ of 0.01 μF/cm^2, while axon nodes had R_m_ of 50 Ω/cm^2^. The models included three voltage-gated conductances: a Na^+^ conductance, a delayed rectifier K^+^ conductance, and two A-type K^+^ conductances. The values of these conductances are assigned according to distance from the soma as described in (Jarsky et al., 2005). While the Na^+^ and 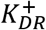 conductances are fixed at 0.04 S/cm2, the value of the 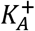 conductances steadily increases from 0.05 S/cm2 at the soma to 0.3 S/cm2 at 500μm from the soma. There is a crossover point between the two different 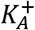 conductances at 100μm from the soma. Furthermore, the extracellular mechanism (Hines and Carnevale, 1997), which allows for injection of extracellular electric potentials was inserted into the models by T2N simultaneously with the other biophysics. Following the generation of the model files by T2N, other necessary files for the next steps are also generated and automatically placed in the correct location.

### FEM Modeling of the TMS induced Electric field

To study the behavior of neurons under non-invasive brain stimulation, the first step is to calculate the electric field generated at the macro- and mesoscopic scale. This includes computing the spatial distribution and time course of the TMS electric field. Since the stimulation frequency is relatively low, we can use the quasi-static approximation to separate the spatial and temporal components of the electric field (Plonsey, 1969, p. 203; Plonsey and Heppner, 1967; Windhoff et al., 2013). For the spatial component, Maxwell’s equations need to be solved for the model of interest. Exact analytical solutions can be determined for simple geometries such as concentric spheres with homogenous electromagnetic properties (Eaton, 1992). However, for more complex geometries such as the human brain, numerical simulations are used to calculate the electric field distribution. Several methods exist to perform these simulations such as the boundary element method (Nummenmaa et al., 2013; Salinas et al., 2009) and the finite element method (Miranda et al., 2003; Wagner et al., 2004; Weiping Wang and Eisenberg, 1994). Here we calculate TMS-induced electric fields using FEM models implemented in the open-source software SimNIBS v3.1 (Saturnino et al., 2019). SimNIBS is a versatile simulation platform that can simulate TMS electric fields for various geometries and a variety of TMS coils.

Under the quasi-static assumption, the time course of the TMS electric field is the same as that of the TMS stimulation output (rate of change (dI/dt) of the coil current). Therefore, after determining the spatial distribution of the electric field, we can find the electric field at any time point by scaling the spatial distribution to the TMS waveform. For further details about the TMS waveform, refer to the ‘Stimulation Waveform Generation’ section below.

### Electric field Coupling to Neuron Models

After calculating the macroscopic TMS electric fields induced in the FEM model of interest, these external fields need to be coupled with the neuron models. In this pipeline, this is performed following: 1) Coordinates of the neuron compartments from the neuron model in the NEURON environment are exported to a text file. 2) The FEM model including the electric fields and the neuron coordinate files are imported to MATLAB. 3) The user enters the desired location and depth (relative to the grey matter surface) of the neuron. 4) Based on the values provided in step 3, the neuron is translated to the desired location. Additionally, the neuron is automatically orientated normal to the grey matter surface as this orientation represents the columnar cytostructure of major neurons (Amunts and Zilles, 2015; DeFelipe et al., 1990; Mountcastle, 1997). However, different preferred orientations can be set if desired. The new neuron compartmental coordinates are calculated based on this coordinate transformation. 5) The electric field at the location of neuronal compartments is interpolated from the macroscopic TMS electric fields calculated in the FEM model. 6) In this step, the user can scale the electric field strength if needed. Since the electric field strength scales linearly with the stimulation intensity, one can easily scale the electric fields instead of rerunning the FEM simulations at different intensities. This allows expediting the simulations e.g. for simulating multiple TMS intensities. Note that this is only true for the stimulus intensity and not applicable if the coil location/orientation, or the FEM model is changed. 7) The quasipotentials are computed over all compartments as described in (Wang et al., 2018). This allows us to convert all necessary information needed to incorporate the external TMS-induced electric fields into a single scalar input at each coordinate of the neuron model. 8) The quasipotentials are written in a file that will be used later in the pipeline for the NEURON simulations. Additionally, the neuron (transformed to the desired location) and the FEM model are exported as mesh files for visualization.

To simplify the multi-scale modeling process, we have also enabled an alternative method to skip the FEM electric field modeling and the corresponding coupling step. In this case, the electric field is assumed to be spatially uniform over the extent of the neuron. This allows the user to specify the TMS-induced electric field everywhere using a single scalar for the amplitude and a vector for orientation. Typically, since neurons are considerably smaller than the TMS coil and the head model, the electric field distribution confined to a single neuron region is mostly uniform. Therefore, the uniform electric field approximation provides sufficiently accurate results in the majority of cases. However, note that the uniform electric field approximation does not always hold. This occurs mainly in the following cases: 1) The neuron crosses a tissue boundary e.g. between Grey matter and white matter (Opitz et al., 2011). Due to the difference in electrical conductivities between tissues, a difference in the electric fields can arise between tissues. 2) The neuron is spatially extended (e.g. neurons with long axonal projections) so that the homogeneity of the electric field over small scales does not apply anymore. 3) The tissue surrounding the neuron is highly inhomogeneous. Although this is a rare scenario since for the purpose of estimating electric fields under non-invasive brain stimulation usually the tissues are assumed to be homogenous in FEM models.

In the case of a uniform electric field, the quasipotentials equation can be simplified to the following expression:

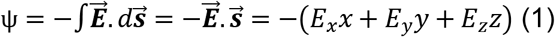

Where 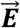 is the electric field, 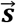 is the displacement vector, E_x_, E_y_, and E_z_ stand for the Cartesian components of the electric field, and x, y, and z denote the Cartesian coordinates of each compartment. Due to the simplicity of this equation, this step is computed in the NEURON environment.

Regardless of whether the electric field is uniform or based on the FEM model, the quasipotentials are calculated at each neuron segments (as exported from the NEURON model) and applied to the neuron simulations by using the extracellular mechanism available in the NEURON environment (Aberra et al., 2018; Hines and Carnevale, 1997). This process accounts for the exogenous fields induced by TMS.

### Stimulation Waveform Generation

As mentioned above, the time course of the TMS electric field follows the first temporal derivative of the stimulation waveform. It is thus very important to accurately represent the TMS waveform to investigate the temporal interaction of the external electric fields with neurons. For repetitive TMS (rTMS) a TMS pulse train is generated based on the parameters of the rTMS protocol. The user has the option to choose the TMS pulse type, inter-pulse interval, and the number of pulses. We included TMS pulse types commonly used in commercial TMS machines i.e. monophasic, and biphasic pulses (Kammer et al., 2001). For the monophasic pulse, we created the waveform based on the equations outlined in (Roth and Basser, 1990). The biphasic pulse was created by using the electrophysiological recording of the TMS pulse induced by MagPro X100 TMS machine (MagVenture, Lucernemarken, Denmark). Based on the specified parameters, the pulses are concatenated to generate a pulse train and then written in a file that is used later in the neuron simulation. Note that advanced users can create custom-waveforms e.g. TBS and cTMS (Peterchev et al., 2010) as long as they follow the same format for existing waveforms.

### Neuron Model Simulations

In this step, the simulation is run based on the generated NEURON model and the files corresponding to the TMS waveform. During this stage, the user is prompted to choose to use the quasipotentials file calculated previously or to proceed with a uniform electric field. In the latter case, the user should enter the intensity of the electric field and its orientation, either in spherical or Cartesian coordinates. Then, after running the simulation, the output files are automatically created. This includes voltage traces of all neuron segments over time and the coordinates of the segments and their connections. If the user intends to continue the pipeline with modeling calcium dynamics, a MATLAB script converts the NEURON results into file formats that are compatible with the next step.

### Calcium Simulations

All necessary components were implemented in the simulation toolbox NeuroBox (Breit et al., 2016). NeuroBox is a simulation toolbox that combines models of electrical and biochemical signaling on one- to three-dimensional computational domains. NeuroBox allows the definition of model equations, typically formulated as ordinary and partial differential equations, of the cellular computational domain and specification of the mathematical discretization methods and solvers (Reiter et al., 2013; Vogel et al., 2013). Built with VRL-Studio (Stepniewski et al., 2019), NeuroBox offers user interface workflow canvases to control the simulation workflow and all biological and numerical parameters. The user can specify simulation parameters for the end time, refinement level, and load the geometry and specify an output location.

#### Calcium Model Equations

Calcium mobility in the cytosol is described by the diffusion equation

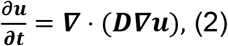

where *u*(*x, t*) is the vector quantity of calcium concentration in the cytosol *[Ca^2+^]* and *calbindin-D28k*. The diffusion constants *D* are defined using data from (4). The interaction between cytosolic calcium and calbindin-D28k are described by

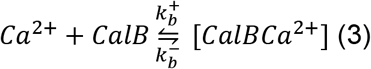

The rate constants 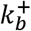 and 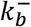 are defined in (Breit et al., 2018). The calcium dynamics are modeled by a system of diffusion-reaction equations on a one-dimensional tree geometry with three spatial coordinates, the equations are as follows:

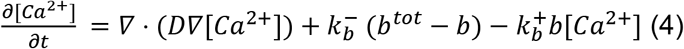

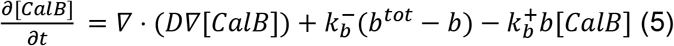

where the concentration of the CalB-Ca^2+^ compound is expressed by the difference of the total concentration of CalB present in the cytosol (*b^tot^*) and free CalB, the former of which is assumed to be constant in space and time (this amounts to the assumption that free calcium and CalB have the same diffusive properties). The parameters used in this study are taken from (Breit et al., 2018).

In order to study the influence of the intracellular organization on Ca^2+^ signals, we include Ca^2+^ exchange mechanisms on the plasma membrane (PM). For the plasma membrane, we consider plasma membrane Ca^2+^ −ATPase pumps (PMCA), Na+/Ca^2+^ exchangers (NCX), calcium release due to voltage-dependent calcium channels (vdcc), and a leakage term. This amounts to the flux equations (number of ions per membrane area and time)

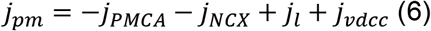

With the Hill equations

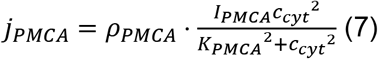

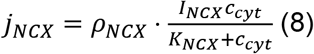

The flux equations for the voltage-dependent calcium channels are given by

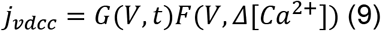

where *G* is the gating function and *F* is the flux function (Borg-Graham, 1999). Both depend on the voltage at the channel at a particular time *t*. For *F, Δ*[*Ca*^2+^]is the difference in the internal and external ion concentration

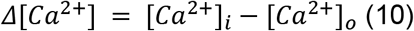

And

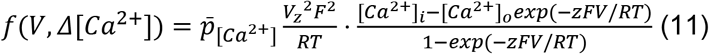

Where R is the gas constant, F is Faraday’s constant, T is in Kelvin, 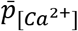 is the permeability of the calcium channel, and z is the valence of the ion (Borg-Graham, 1999).

The gating function *g* is described by a finite product

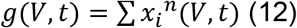

Where *x_i_* is the open probability of the gating particle, in this case, it is only calcium, and *n* is the number of particles. The open probability is described by the ODE

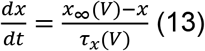

Where *x*_∞_ is the steady-state value of x, and *τ_x_* is the time constant for the particular particle x, formulas are given in (Borg-Graham, 1999).

#### Numerical Methods for Calcium Simulations

For numerical simulations, the equations are discretized in space using a finite volumes method. Current densities, across the plasma membranes, can be incorporated into the reaction-diffusion process very naturally and easily this way. Time discretization is realized using a backward Euler scheme, i.e., for each point in time t, the term 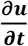 is approximated by

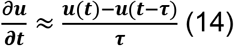

Where *τ* is the time step size. For the results we present here, the emerging linearized problems were solved using a Bi-CGSTAB (Breit et al., 2018) linear solver preconditioned by an incomplete LU decomposition.

### Visualization

Additionally, we have provided a sample script that can generate a video visualizing the 3D distribution of the membrane potentials and the calcium concentrations based on the simulated data from the previous steps. In this procedure, the snapshot of voltage/calcium spatial distribution at each time step is displayed in Gmsh (Geuzaine and Remacle, 2009) and then captured as a video frame. In the end, by concatenating these frames together, a video is created. This script is capable of visualizing the voltage traces and calcium concentrations separately or next to each other in a single video file for easier comparison. Alternatively, users can visualize the data with Paraview (Ahrens et al., 2005).

## Acknowledgments

We thank Swathi Anil, Dr. Zsolt Turi, and Dr. Harry Tran for the helpful discussions and testing of the toolbox. This work was supported by the National Institutes of Health (R01MH118930 to GQ and AO and R01NS109498 to AV and AO and RF1MH117428 to AO), Federal Ministry of Education and Research Germany (BMBF, 01GQ1804A to AV; BMBF, 031L0229 to PJ), and the von Behring Röntgen Foundation (to PJ).

