## Supplementary-materials for "Multi-scale Modeling Toolbox for Single Neuron and Subcellular Activity under (repetitive) Transcranial Magnetic Stimulation": Sup-Multi-scale.pdf

### Supplementary Data

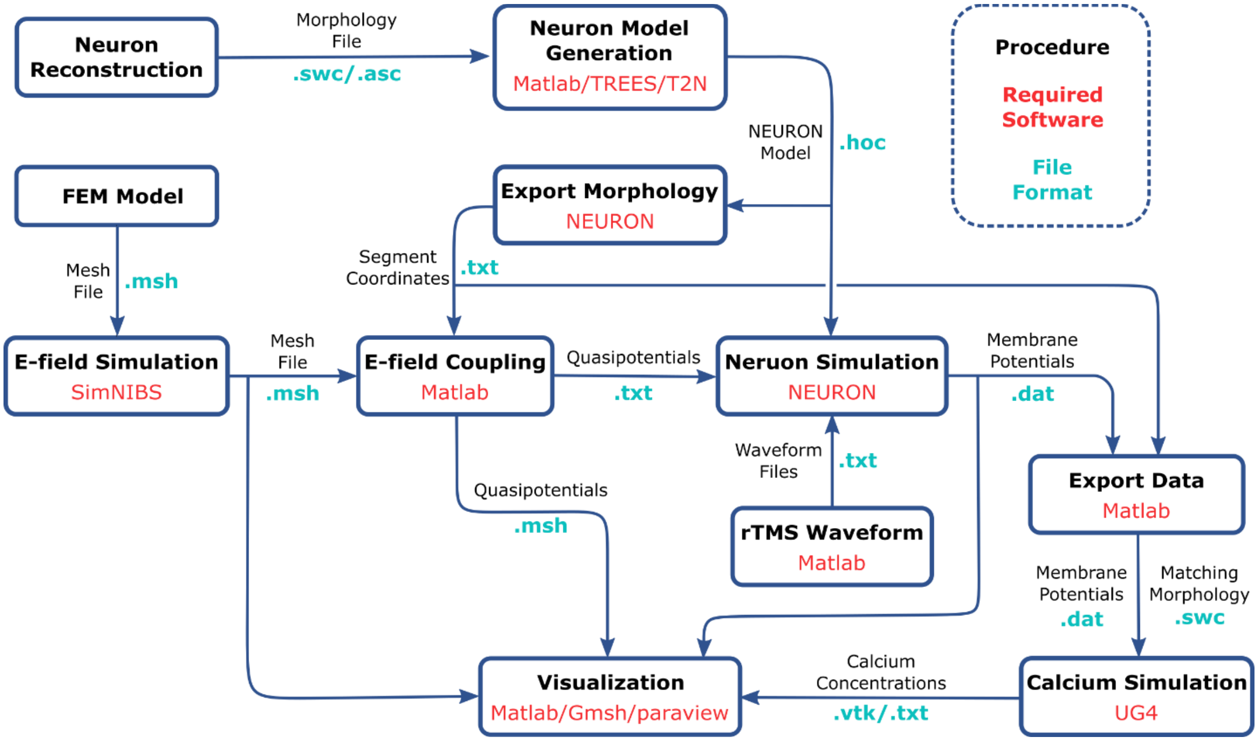


**Figure S1.** Pipeline overview. Each step of the procedure is represented in a box. The software used for each step are shown in red. The file formats of the input and output files are shown in blue.

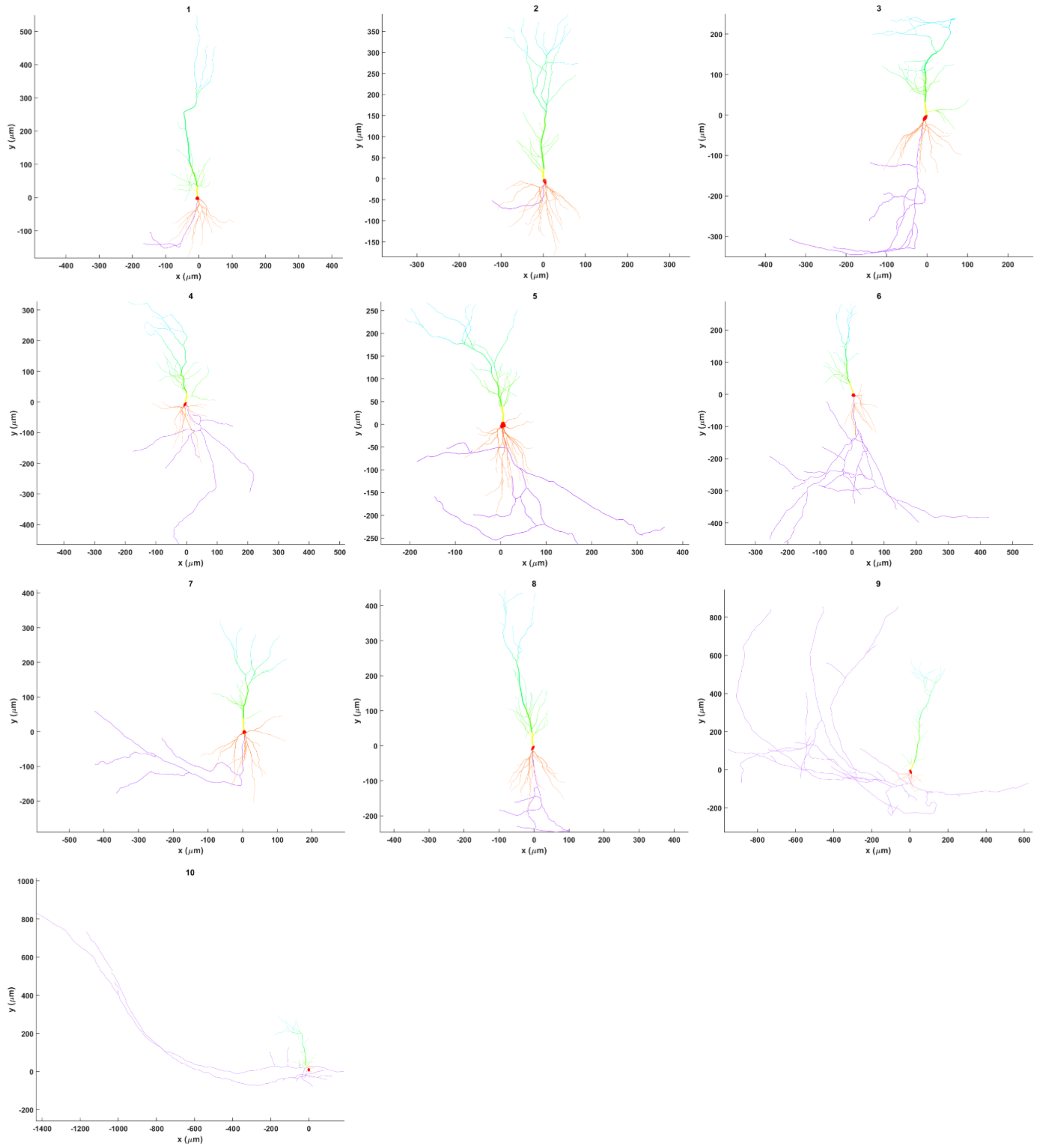


**Figure S2.** Morphological reconstruction of the 10 sample CA1 pyramidal neurons. Red, orange, yellow, green, blue, and purple branches respectively represent soma, basal dendrites, proximal apical, distal apical, apical tufts, and axon.
